# Genomic diversity and novel genome-wide association with fruit morphology in Capsicum, from 746k polymorphic sites

**DOI:** 10.1101/487165

**Authors:** Vincenza Colonna, Nunzio D’Agostino, Erik Garrison, Jonas Meisner, Anders Albrechtsen, Angelo Facchiano, Teodoro Cardi, Pasquale Tripodi

## Abstract

**Background:** *Capsicum* is one of the major vegetable crops grown world-wide. Current subdivision in clades and species is based on morphological traits and coarse sets of genetic markers. Fruits broad variability has been driven by breeding programs and has been mainly studied by linkage analysis.

**Results:** We discovered 746k variable sites by sequencing 1.8% of the genome in a collection of 373 accessions belonging to 11 *Capsicum* species from 51 countries. We describe genomic variation at population-level, confirm major subdivision in clades and species, and show that the known subdivision of *C. annuum* in two groups separates large and bulky fruits form small ones. In *C. annuum,* we identify four novel loci associated with phenotypes determining the fruit shape, including a non-synonymous mutation in the gene *Longifolia 1-like* (CA03g16080).

**Conclusions:** Our collection covers all the economically important species of *Capsicum* widely used in breeding programs, and represent the widest and largest study so far in terms of the number of species and genetic variants analyzed. We identified a large set of markers that can be used for population genetic studies and genetic association analyses. Our results foster fine genetic association studies and foresee genomic variability at population-level.

## Background

Pepper (*Capsicum* spp.) is one of the major vegetable crops grown worldwide, largely appreciated for its economic importance and nutritional value. Pepper originated in North-West and South America and then expanded and diversified in the other Southern and Central American regions. The taxonomic arrangement of the genus have been recently revised establishing the existence of 35 species distributed in 11 clades,[1] the most important of which are Annuum (*C. annuum, C. frutescens, C. chinense*) Baccatum (*C. baccatum, C. praetermissum, C. chacoense*) and Pubescens (*C. pubescens*). Population structure of *Capsicum* spp. inferred from genetic markers mostly reflects the known taxonomic grouping, however it highlights sub-groups within *C. annuum* and admixture between *C. chinense* and *C. frutescens.*[2, 3] Even if these studies were carried out on a large number of accessions, both are based on small number of markers (28 simple sequence repeats, SSR, on 1,400 accessions and 48 simple nucleotide polymorphisms, SNP, on 3,800 accessions), therefore the population structure of the different *Capsicum* species has never been investigated deeply.

Pepper presents a wide diversity in fruit size and shape achieved after a successful selection process. The early steps of domestication involved key traits such as non-deciduous fruits and the orientation of fruit tip from erect to pendant. Further steps of selection resulted in wide variability of fruit shape and size comparable to other *Solanaceae.*[4] Nevertheless, unlike other cultivated *Solanaceae,* e.g. tomato,[5, 6, 7, 8] the genetics underlying fruit shape in pepper is limited to findings from quantitative trait *loci* (QTL) analysis in a few biparental mapping populations (Supplementary Table 1), mostly phenotyped with low-throughput techniques,[9, 10, 11, 12, 13, 14, 15, 16] and/or limited to the identification of QTLs.[17, 18, 19, 20] As a consequence, major genetic *loci* underlying *Capsicum* fruit variability have not yet been identified.

Technical advances in high-throughput phenotyping allow to precisely measure the determinants of fruit shape and size using standard ontologies,[21] while the availability of a reference sequence[22] and population-based sequence data enables the discovery and fine mapping of genomic variants. The combination of these two factors lays the foundation for a better investigation of the genetics of fruit size and shape through genome-wide association studies (GWAS), as carried out for capsacinoid content[23, 24] and peduncle length[25], but never performed for fruit morphology attributes in *Capsicum*.

In this study we discovered 746k high-quality polymorphic sites from analyzing sequence data of 373 pepper accessions from eleven species of *Capsicum* and measured thirty-eight fruit shape and size attributes in 220 *C. annuum* accessions. We used these data to (i) uncover genomic properties of the pepper genome, (ii) describe population structure within the *Capsicum* genus at a resolution never achieved before, (iii) study natural selection, and (iv) discover significant association between genetic markers and traits related to pepper fruit shape and size in *C. annuum*. Finally, we discovered a non-synonymous change in the sequence of *Longifolia 1-like* gene associated with variance in *C. annuum* fruit elongation.

## Results

### Genomic diversity of the *Capsicum* genus at 746k variable sites

The germplasm collection presented here covers all the economically important species of *Capsicum* widely used in breeding programs. It includes 373 accessions belonging to eleven *Capsicum* species, of which five are wild (Table 1). Two hundred and twenty *C. annuum* accessions were already described.[26] With the exception of 48 accessions for which the geographical origin is unknown, the remaining accessions are from 51 countries (Supplementary Table 2, Figure 1).

**Table 1.** Summary of the samples used in this study. A species was included in a specific analysis based on the number of accessions available. Horizontal lines mark clusters of species used in different analyses.

| Species | # of diploid accessions | Type |
| --- | --- | --- |
| *C. annuum* | 220 | domesticated |
| *C. chinense* | 62 | domesticated |
| *C. baccatum* (*varpendulum*) | 39 | domesticated |
| *C. frutescens* | 14 | domesticated |
| *C. chacoense* | 13 | wild |
| *C. pubescens* | 13 | domesticated |
| *C. annuum* var *glabriusculum* | 5 | wild |
| *C. praetermissum* | 3 | wild |
| *C. baccatum* var *baccatum* | 2 | domesticated |
| *C. galapagoense* | 1 | wild |
| *C. eximium* | 1 | wild |

**Figure 1.**
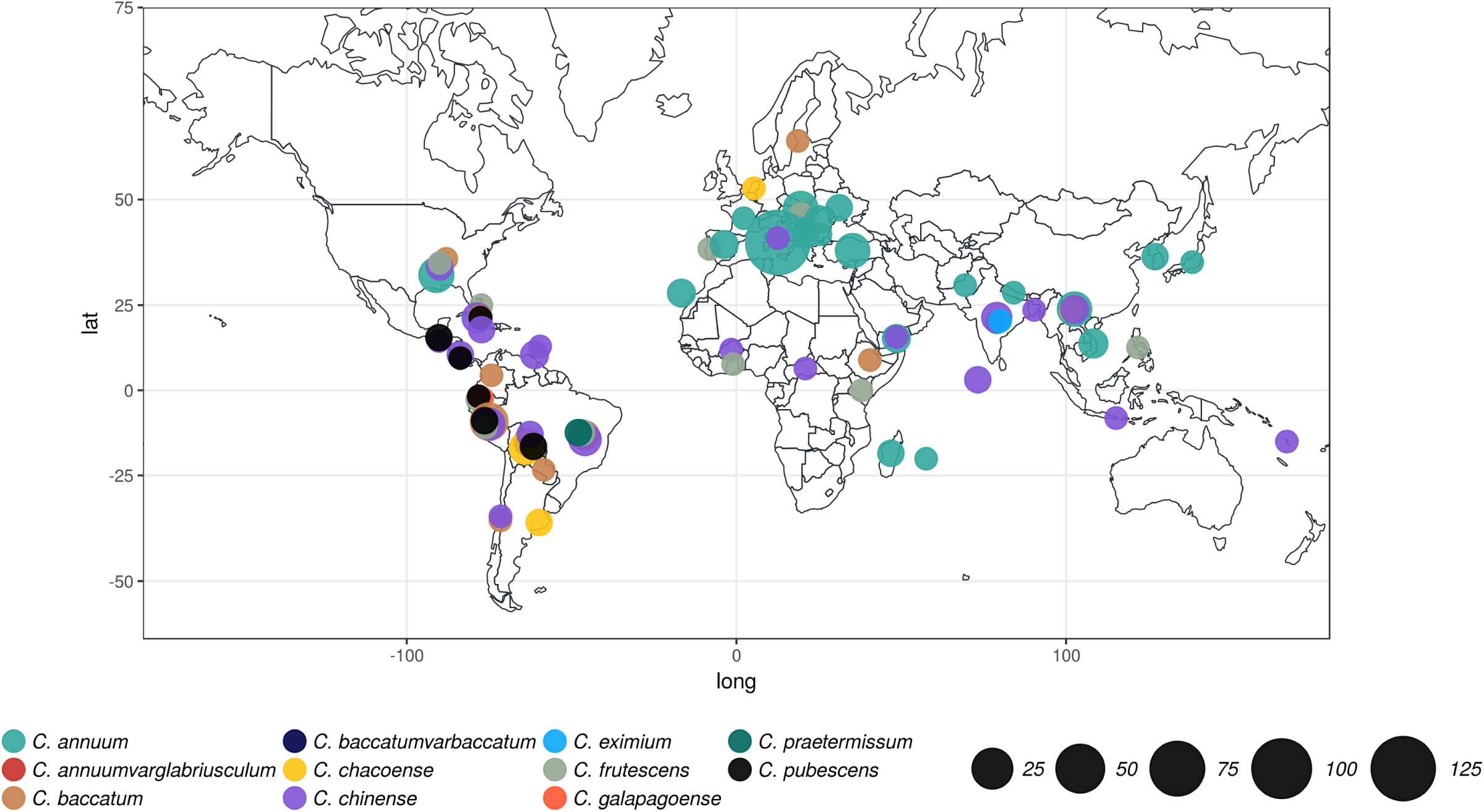
Geographical origin of the Capsicum accessions presented in this study. With the exception of 48 accessions deriving from the germplasm bank for which the origin is unknown. Circle colors define species while size is proportional to sample size.

Genomic DNA extracted from accessions was digested with the restriction enzyme *Ape*KI obtaining more than 7.5M master tags[26] that were assembled in 605,076 genomic regions or fragments. The cumulative sequence length of the fragments is 48,869,949 bp, corresponding to 1.8% of the genome, with average depth-of-coverage 5.8. Fragments are scattered across the genome. Average fragment size is 81 bp (standard deviation is 42.7 bp), and the range of distance between two consecutive fragments varies from 3 to 278,100 bp (average 4,551 bp). The majority of the fragments (85.9%) are intergenic.

By aligning sequence reads along the reference genome,[22] we identified 1,382,545 polymorphic sites, of which 746,450 have Phred-scaled quality scores (QUAL) >10 and were considered for all downstream analyses (Figure 2a). These figures compared to the cumulative sequence length of the fragments suggest that 1.5% of the genome is variable in *Capsicum.* Up to 95% of variants are single nucleotide polymorphisms (SNP, 82.4%) or multi-nuclotide polymorphisms (MNP, 12.5%). Insertions and deletions (InDels) represent 1.9% of all variants and their size ranges from deletions of 30 nucleotides to insertions of 20 nucleotides. InDels of three or multiples of three nucleotides are more frequent in genic region compared to intergenic ones, suggesting a preference for InDels that add or remove triplets over those causing frame-shifts mutations (Supplementary Figure S1). Variants are not equally distributed in the genome, and in fact, only 37.1% of the fragments have at least one variant. Average number of variants per 100 bp is 9.4 and 12.3 in genic and intergenic regions respectively (Figure 2b) and this difference is significant (Mann-Whitney test p-value <2×10e-16). Finally, based on the annotations,[22] only 5.22% of variants (38,964) fall within exons.

**Figure 2.**
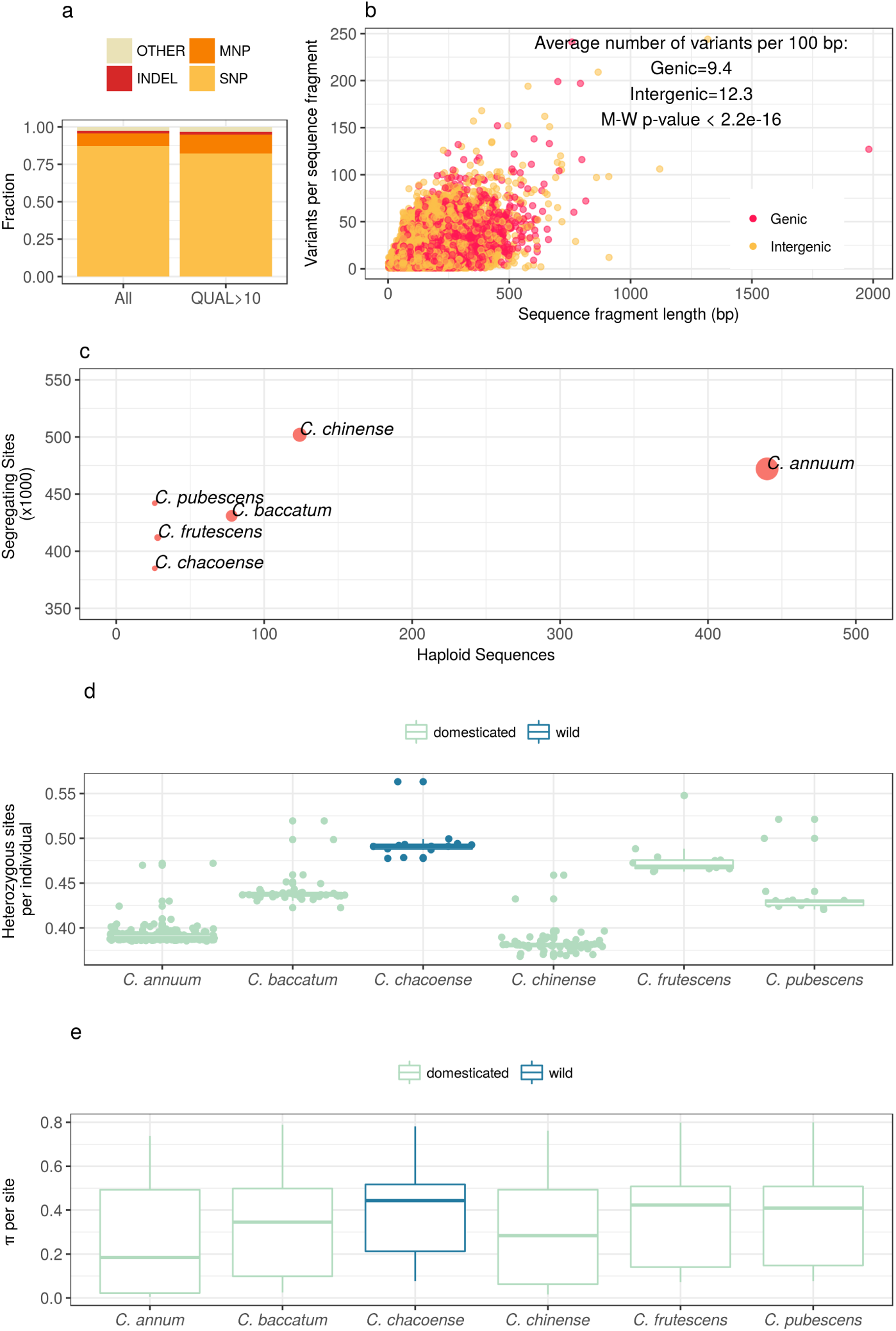
Genomic diversity in Capsicum. **(a)** Types and abundance of variant types. The majority of variants is single nucleotide polymorphisms (SNP), followed multi-nucleotide polymorphisms (MNP) and insertions/deletion (INDEL). A very small fraction of variants is complex combinations of SNP, MNP, and INDEL. QUAL>10 refers to Phred-scaled quality scores. **(b)** The number of variants per sequence fragment normalized by the fragment lenght. Intergenic sequences have higher number of variants, suggesting that intergenic regions are less constrained on variation. Each circles is a sequence fragments and colors distinguish genic from intergenic ones. **(c)** Average number of segregating sites per species is 440.6k. The number of segregating sites per species is roughly proportional to sample size with the exception of *C. annuum* for which there are less variable sites than expected given the number of accessions, most likely because of intensive domestication. **(d)** Proportion of heterozygous sites per accession. Species that underwent extensive domestication (*C. annuum* and *C. chinense*) have very low heterozygosity, while the wild species *C chacoense* has the highest variability. **(e)** Nucleotide diversity per site (*π*) follows the same trend of the heterozygosity.

For six of the eleven *Capsicum* species with >10 accessions, we further investigated genomic diversity. The average number of segregating sites *per* species is 440,600 and it is roughly proportional to sample size. *C. annuum* is an exception, as it seems less variable than expected given the number of haploid sequences, which likely reflects the recent loss of diversity due to intensive selective breeding (Figure 2c). In fact, *C. annuum* accessions are among the least heterozygous (Figure 2d) and also those with the lowest nucleotide diversity (Figure 2e). By contrast, the wild species *C. chacoense* has the highest values of heterozygosity per individual (Figure 2d) and nucleotide diversity per site (π, Figure 2e), even having the lowest number of segregating sites compared to domesticated species with a similar number of haploid sequences (Figure 2c).

To summarize, we discover 746k variable sites by analyzing 605k fragments of average size 81 bp that cover 1.8% of the genome of 373 *Capsicum* accessions. Variant density is not uniform, and in fact, only 37.1% of the fragments contain variants. Variant density is significantly lower in genic regions compared with intergenic ones. The majority of variants are single nucleotide changes. Due to our use of Genotype By Sequencing (GBS) and reference-guided analysis, we were only able to discover InDels up to a few tens of bp. Among species, *C. annuum* is the least diverse while *C. chacoense* is the most variable, despite possible variant detection bias due to the fact that the reference sequence belongs to *C. annuum*.

### Population structure of *Capsicum* reveals strong subdivisions with little or no admixture among species

We investigated the population structure of the nine *Capsicum* species with at least two accessions (Table 1) using data from the 746k variable sites and three approaches. Phylogenetic reconstruction confirms clustering of accessions in species (Figure 3a), as observed in similar studies based on plant morphological characteristics[27, 28] or a smaller number of genetic markers.[3, 29, 2] Nevertheless, this is to date the largest phylogenetic study in terms of genomic markers analyzed in *Capsicum* species. We used principal components analysis (PCA) to summarize the observed genetic variation among accessions and species. The first two components explain the 24% of the observed variation separating three main domesticated species: *C. annuum, C. baccatum,* and *C. chinense.* While describing almost one-fourth of the variance, clustering within the first two components is not complete, and a number of accessions are positioned in between clusters. The third and fourth components (5% of explained variance) separate *C. pubescens* and *C. chacoense* between them and from the cluster of domesticated species (Figure 3b). While *C. chacoense* is a wild species, *C. pubescens* is domesticated, although one of the least easy to breed.[30] PCA clustering is not related to geography (Supplementary Figure S2).

**Figure 3.**
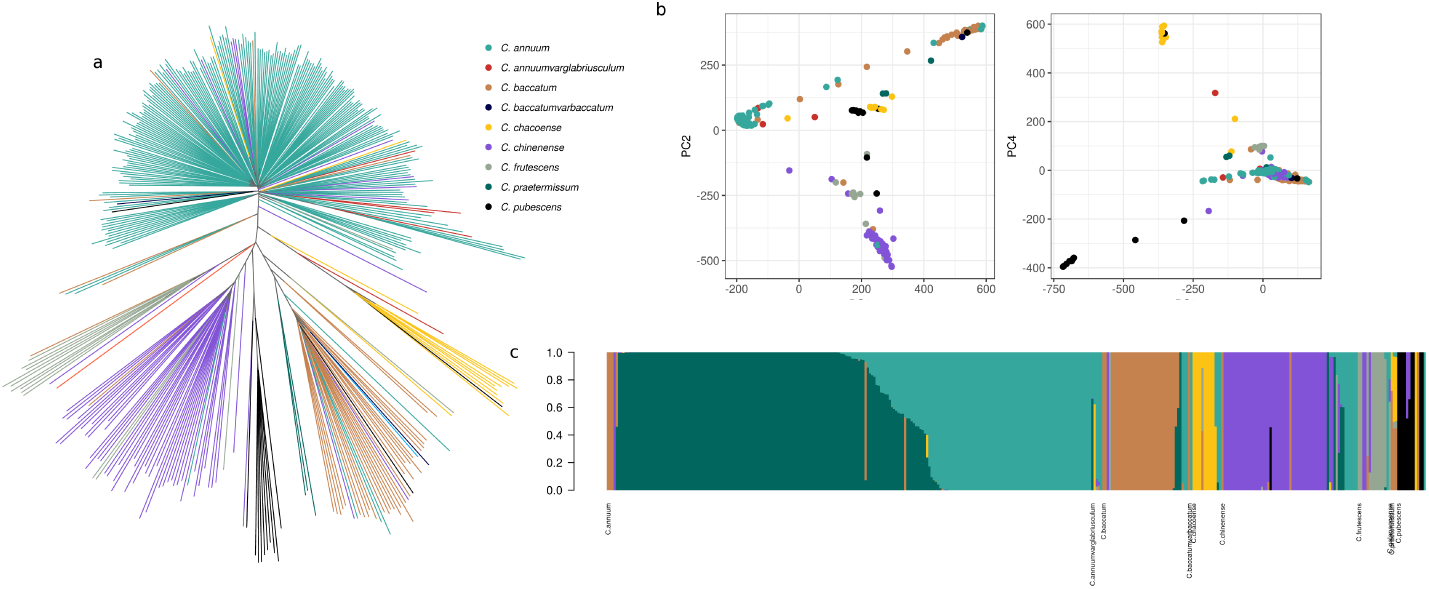
Population structure of the Capsicum species derived form 746k genomic variants. **(a)** Phylogenetic reconstruction of the relationships between the accessions. With a few exceptions, clusters correspond to species. **(b)** Principal component analysis. The first two components separate the three main domesticated species. Clustering within the first two components is not complete, and a number of accessions are positioned in between clusters. The third and the fourth components separate *C. pubescens* and *C. chacoense* between them and from the cluster of domesticated species. (**c**) Model based admixture analysis in the hypothesis of seven clusters. With the exceptions of few admixed or misplaced individuals, clusters correspond to species and within *C. annuum* is possible to observe two groups with distinct genetic features.

Finally, admixture analysis supports subdivision into seven clusters (Figure 3c, Supplementary Figure S3). Clusters correspond mostly to species, with the exception of the further subdivision of *C. annuum* into two sub-clusters with some admixed individuals, as observed in other studies.[26, 3, 2] Most accessions belong to only one cluster, with the median coefficient of membership to the best-matching cluster being 0.99 (mean ±sd is 0.9471 ±0.13, Supplementary Figure S4). Nevertheless, some of the accessions seem misplaced probably because of mislabeling or misclassification[31] by germplasm providers. Like PCA, admixture clustering is not related to geography (Supplementary Figure S5).

In conclusion, the population structure analysis presented here shows that accessions cluster according to species without regards to their geographical origin and according to the known phylogenetic relationships. There is little or total lack of admixture among species and instead, deep clustering analysis reveals a major subdivision within *C. annuum,* as observed before.[26, 3, 2]

### Insight into *C. annuum* fruit variability from high resolution phenotyping

Fruit shape and size varies widely between pepper accessions. We carried out phenol-typing of the 220 accessions of *C. annuum* at thirty-eight phenotypic traits (Figure 4), in order to describe and understand the range of variation of *C. annum* fruits. For each accession, each trait value is the average of forty-eight measures: accessions were grown in triplicate and for each triplicate eight fruit were considered, for a total of twenty-four fruits per accession. Fruits were longitudinally sectioned in half and each section measured. Trait values were scored from the images obtained with a CanoScan LiDE. Measured traits can be classified into two broad categories according to if they are related to fruit shape or size, and the shape category is further organized in eight classes (Supplementary Table S3).

**Figure 4.**
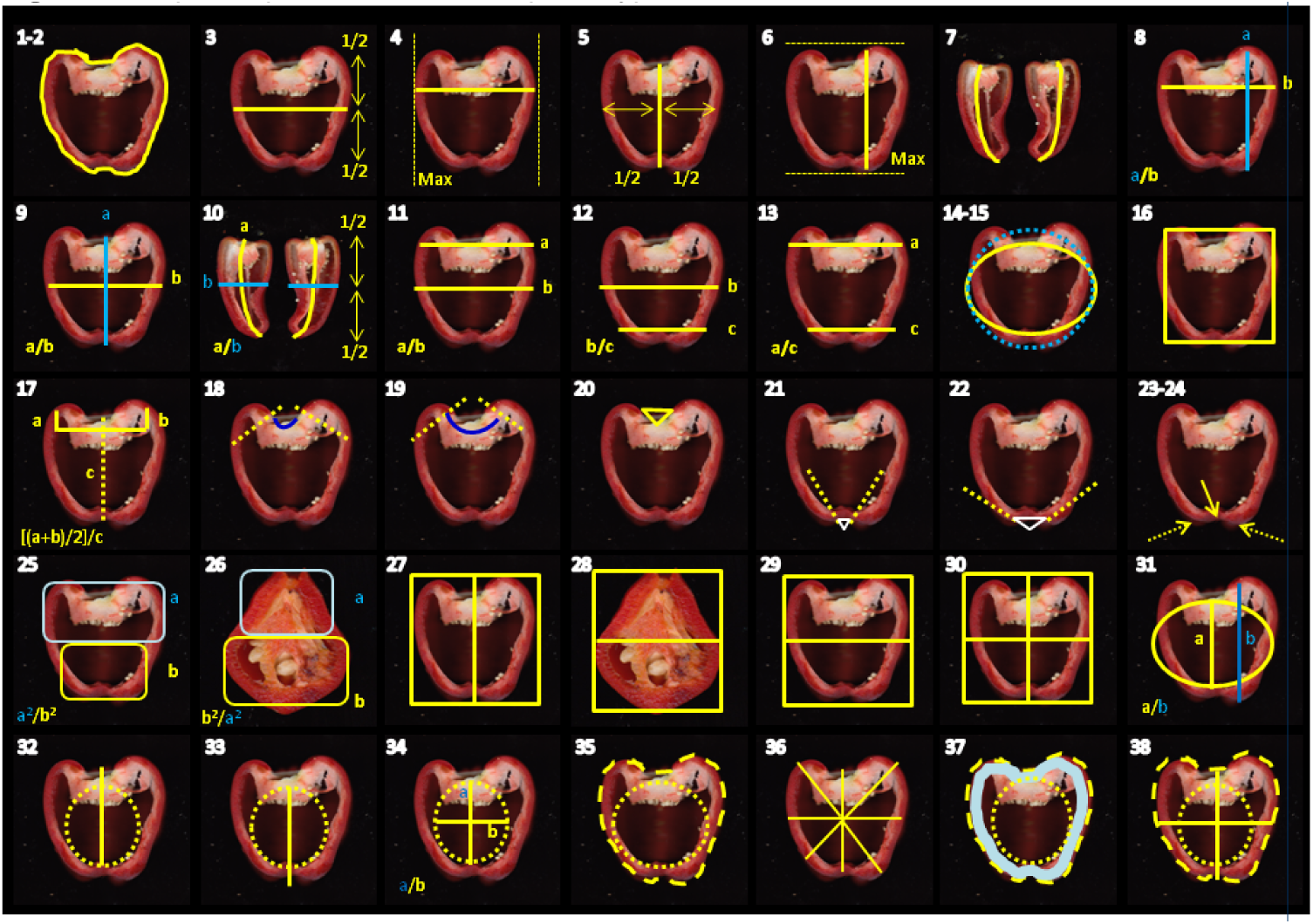
Graphic representation of the phenotypes measured. The numbers refers to Table S3

Traits display a wide variability relative to each other and in fact the average coefficient of variation (CV) is 61% (median CV= 45%) and the largest CV is >300%, suggesting that in many cases the standard deviation exceeds the mean value of the trait (Figure 5a). The less variable trait is the Pericarp Area (CV=7%), i.e. the ratio of the area within the pericarp boundary to the area of the fruit, suggesting that fruits tend to have comparable skin thickness. Other traits with CV<10% are ProximalEccentricity, Eccentricity, and PericarpThickness, all in fact related to the thickness of the pericarp and mostly related to fruit latitudinal section features and eccentricity (Supplementary Table S3). By contrast, the most variable trait is the HAsimmetryOb (CV=346%), which represents the fruit length as the average distance between a horizontal line through the fruit at mid-height and the midpoint of the fruit’s height at each width. Other traits with CV >100 are VAsymmetry, Obovoid, and DistalIndentationArea, all defining the fruit shape (Supplementary Table S3). Pairwise Spearman’s rank correlations among all possible pairs of the 38 traits are statistically significant with p-value ≥10e-7 in 48.9% of the cases (Supplementary Table S4) and it is possible to observe clusters of correlation pairs (Figure 5b). While fruit thickness does not show significant variability, the major phenotypic variation involves traits determining the fruit asymmetry and the distal fruit-end shape.

**Figure 5.**
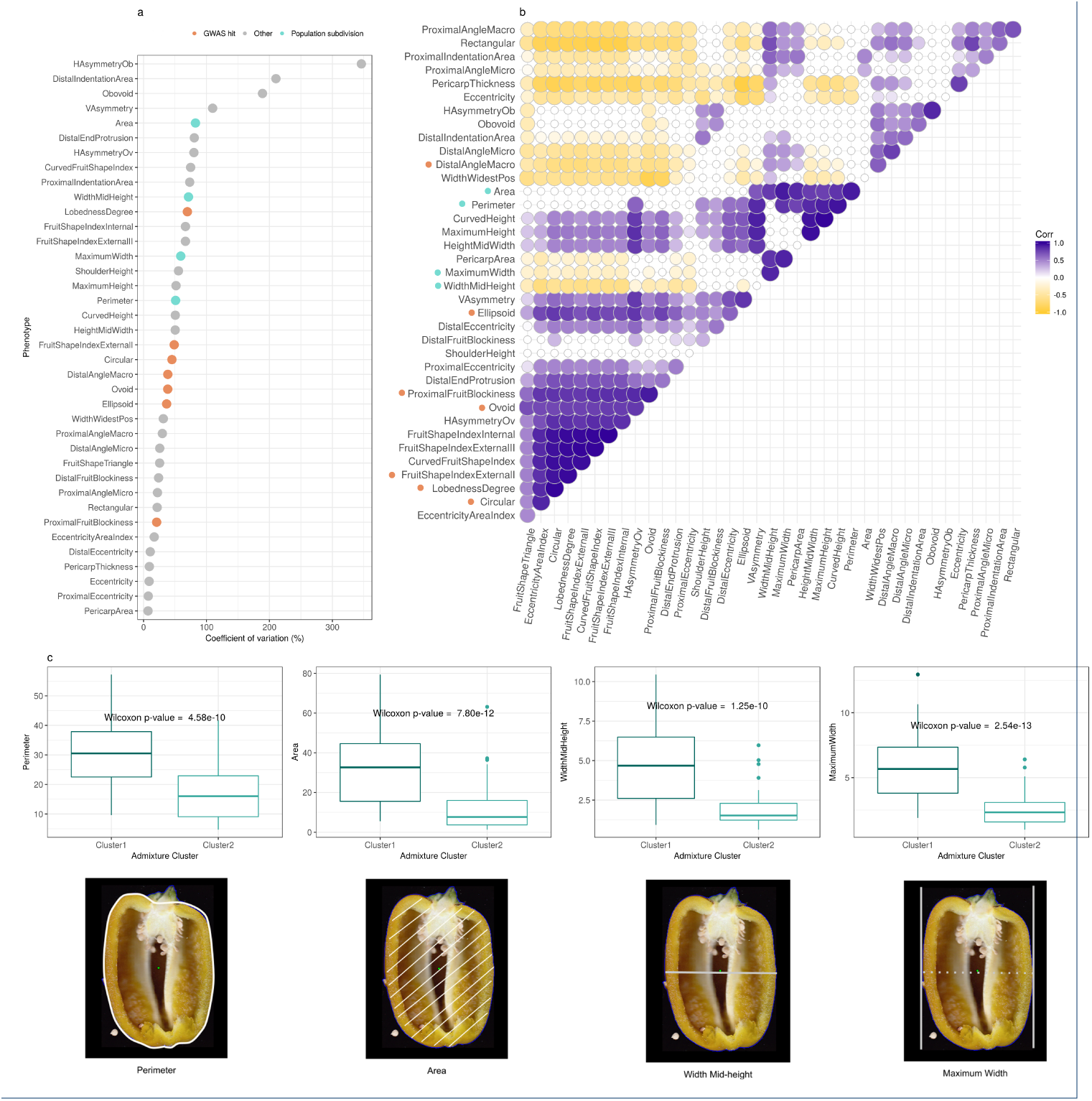
Analyses of thirty-eight quantitative traits related to fruit shape and size. Description of phenotypes are available in Supplementary Table S3. **(a)** Coefficients of variation (CVs) show that very often the standard deviation exceeds the mean value of the trait, suggesting a great variability of the traits. **(b)** Spearman’s rank correlation coefficients between pairs of phenotypes. Only correlation coefficients with p-value <10e-7 are shown. In **(a)** and **(b)** green dots mark phenotypes that are significantly different between clusters of *C. annuum* identified in the admixture analysis, while orange dots mark phenotypes showing significant association with genetic markers in genome-wide association tests. **(c)** Traits that significantly differs between the two subgroup of *C. annuum* identified from genetic clustering analysis. Cluster 1 contains bulkier and larger fruits compared to Cluster2.

Subdivision of *C. annuum* accessions into two clusters (Figure 3c) is associated with significant differences in traits related to fruit size (Figure 5c). To remove admixed accessions, samples from *C. annuum* were assigned to genetic clusters determined in the admixture analysis if membership to the cluster was >90%. Of the 38 traits, Area, MaximumWidth, WidthMidHeight, and Perimeter are significantly different between the two genetic clusters (Figure 5c). With the exception of the pair Perimeter-WidthMidHeight, all other pairs among the four traits are positively and significantly correlated, and their CV range from 50% (Perimeter) to 82% (Area, Figure 5a,b). Fruit size is significantly different between the two genetic clusters, with Cluster 1 having bulkier and larger fruits relative to Cluster 2. This suggests that differences in fruit size in *C. annuum* might be caused by genetic differences.

### Genetics of fruit shape in *C. annuum*

Having observed concordance between genetic and phenotypic clustering in *C. annuum* we next aimed at understanding the genetic basis underlying the natural variation in fruit size. We carried out genome-wide association analyses between the thirty-eight traits and the 746k genetic markers identified in this study using a univariate linear mixed model. To account for cryptic population structure and relatedness we integrated in the model a relatedness matrix estimated from the accession genotypes as random effects. In addition, we included the first two PCs (35.7% of explained variance) as fixed effects. SNPs were filtered for missingness (< 5%), minor allele frequency (>1%), and Hardy-Weinberg equilibrium (p-values >0.001), leaving 559,684 SNPs for the association analyses.

Association tests identified eight variants on three chromosomes, significantly associated (Bonferroni corrected p-value <1.78e-08) with seven traits (Table 2, Figure 6a, Supplementary Figure S6). All the seven traits are highly and significantly correlated (Figure 5b) and contribute to determine whether a fruit is circular or elongated (Supplementary Table S3).

**Table 2:** Loci significantly associated with phenotypes related to fruit shape with genome-wide significance. Min/Maj = minor and major alleles; MAF = Minor Allele Frequency; (*β* = Coefficient describing the effect size of the marker in the univariate linear model of association; SE = Standard Error of *β*; p-value = genome-wide Bonferroni corrected p-value for association

| Chr:Position | Genomic region | Nearest gene (kb) | Min/Maj alleles | MAF | Phenotype | $\beta$ | SE | p-value |
| --- | --- | --- | --- | --- | --- | --- | --- | --- |
| 3:183386147 | CA03g16080 | 0 | C/T | 0.13 | Circular<br>Ellipsoid<br>Fruit Shape Index External I<br>Lobedness Degree | 0.10<br>0.03<br>0.91<br>16.97 | 0.016<br>0.005<br>0.145<br>2.449 | 1.36E-08<br>1.58E-10<br>4.93E-09<br>1.38E-10 |
| 10:28759675 | Intergenic | CA10g04730 (-911) | T/C | 0.36 | Distal Angle Macro | 18.47 | 2.962 | 4.97E-09 |
| 10:28759685 | Intergenic | CA10g0473 (-910) | T/G | 0.36 | Distal Angle Macro | 18.47 | 2.962 | 4.97E-09 |
| 10:33533810 | Intergenic | CA10g05040 (166) CA10g05050 (-156) | C/T | 0.22 | Ovoid<br>Proximal Fruit Blockiness | -0.08<br>-0.12 | 0.012<br>0.018 | 8.87E-10<br>1.91E-10 |
| 10:33533831 | Intergenic | CA10g05040 (167) CA10g05050 (156) | T/A | 0.22 | Ovoid<br>Proximal Fruit Blockiness | -0.08<br>-0.12 | 0.012<br>0.018 | 8.87E-10<br>1.91E-10 |
| 10:33534094 | Intergenic | CA10g05040 (169) CA10g05050 (-156) | A/G | 0.22 | Ovoid<br>Proximal Fruit Blockiness | -0.08<br>-0.12 | 0.012<br>0.018 | 8.87E-10<br>1.91E-10 |
| 10:33557960 | Intergenic | CA10g05040 (408) CA10g05050 (-132) | C/G | 0.1 | Ovoid<br>Proximal Fruit Blockiness | -0.15<br>-0.24 | 0.024<br>0.036 | 5.43E-09<br>4.04E-10 |
| 11:18204293 | Intergenic | CA11g04570 (849) CA11g04580 (-892) | T/A | 0.06 | Ellipsoid | 0.04 | 0.006 | 4.39E-09 |

**Figure 6.**
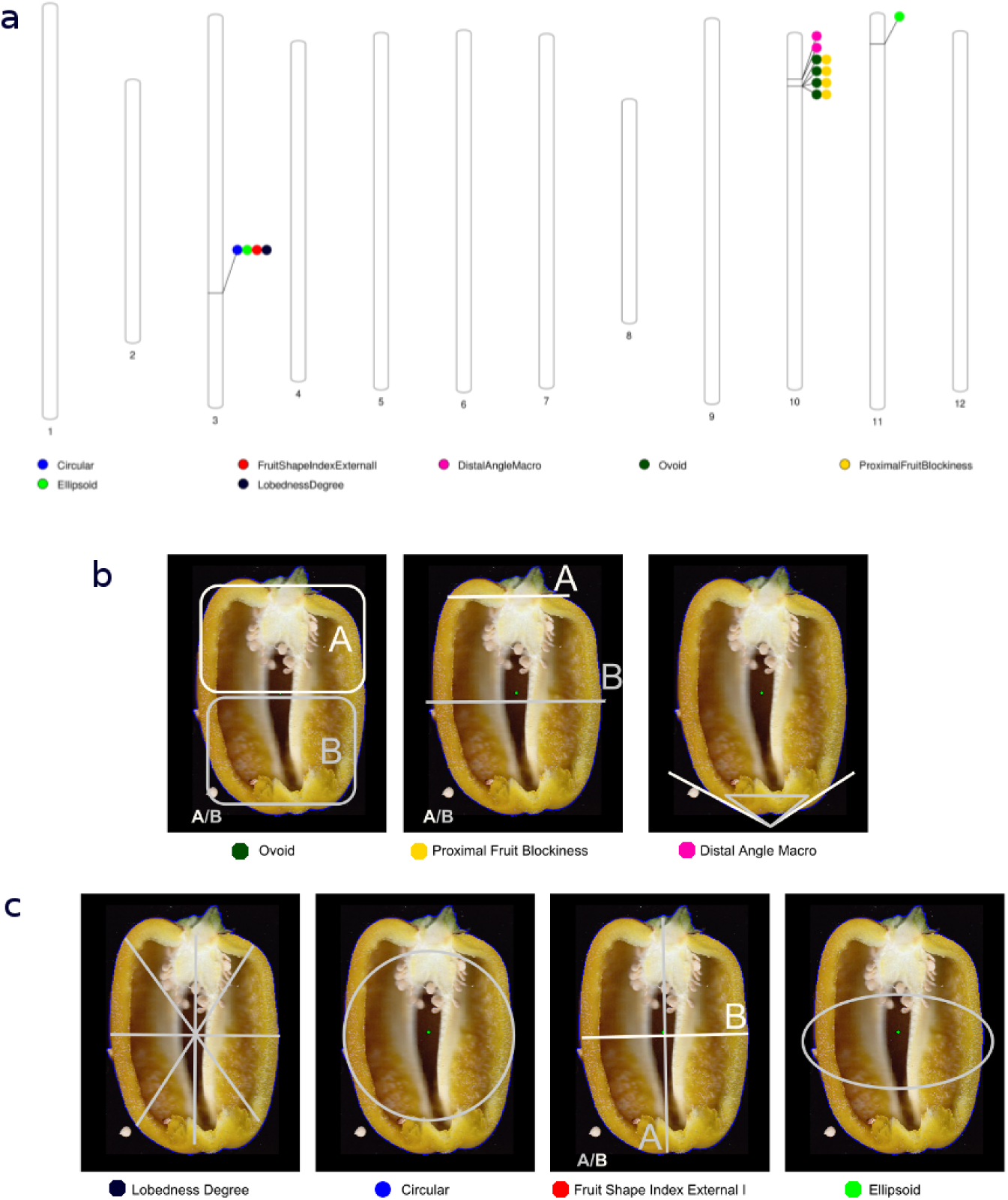
Results of the genome-wide association analysis. **(a)** We identified eight variants at four loci on three chromosomes, significantly associated with seven traits. Circles represent association between one genetic variant and one trait. On chromosome 10, variants are adjacent. Colors distinguish phenotypes. **(b)** Cluster of phenotypes determining whether fruits are pointed or squared. **(c)** Cluster of phenotypes determining if fruits are circular or elongated, with significant association with a variant causing a non-synonymous mutation in the gene *Longifolia 1-like* on chromosome 3.

On chromosome 10 we observe two clusters of variants separated by 4.7 Mbp associated with traits determining whether a fruit is squared or pointed through evaluation of asymmetry, blockiness and the shape of the distal end of the fruit (Figure 6b). The first cluster includes four intergenic variants in a 24 kb region (three of them are in complete linkage) associated with the traits Ovoid and ProximalFruitBlockiness. Both traits specify to what extent a fruit is squared. Ovoid is quantified as the ratio of the area of the fruit above and below middle height (aA/aB Figure 6b), while the ProximalFruitBlockiness is the ratio of the width at the upper blockiness position to width mid-height (wA/wB Figure 6b). In both cases, values close to 1 indicate a rather square fruit, while values >1 are typical of pointed fruits. For the strongest associated SNP 10:33557960 each minor allele confers a more pointed fruit by reducing the ratio for both Ovoid (*β*=-0.15 aA/aB, P=5.43E-09) and for ProximalFruitBlookiness (*β*=-0.24 wA/wB, P=4.04E-10).

The second cluster on chromosome 10 includes two intergenic variants 10 bp apart and in full linkage, associated with the trait DistalAngleMaoro that reflects how much a pepper is pointed and measures the angle between best-fit lines drawn through the fruit perimeter on either side of the distal end point: the smaller the angle the more pointed the pepper. Each minor allele for variants 10:28759675 and 10:28759675 increases is estimate to increase the angle by 18° (*β*=18.47°, *SE* = 2.96, P=4.97E-09).

On chromosome 11 we detect a significant association between one intergenic SNP and the Ellipsoid trait. The Ellipsoid index fits precision of the actual shape of the fruit to an ellipse: smaller values indicate ellipsoidal fruits and measures the fraction of area left out of a fitted ellipse (a.o.e.). The range of Ellipsoid in our data vary from 0.002 a.o.e. to 0.21 a.o.e., and the effect of each minor allele increases of the a.o.e.(*β*=0.04 a.o.e., *SE* = 0.006, P=4.39E-09), contributing to produce pointed fruit.

On chromosome 3, the SNP 3:183386147 is significantly associated with four traits also related to fruit shape: Circular, Ellipsoid, FruitShapeIndexExternalI, and LobednessDegree (Figure 6c, Table 2). Similar to Ellipsoid, Circular fits precision of the actual shape of the fruit to a circle: smaller values indicate circular fruits and measures the fraction of area left out of a fitted circle (a.o.c.). The range of values for Circular in our data set is 0.04-0.46 a.o.c. and each minor allele increase the a.o.c. (*β*=0.10 a.o.c., *SE* = 0.01, P=1.36E-08), as well as the a.o.e. (*β* = 0.03, *SE* = 0.005, P=1.58E-10) contributing therefore to make the fruit elongated. Consistently, the minor allele increases the LobednessDegree, i.e. the dispersion of several distances from the center of weight to the perimeter, up to 16.97mm (*β* = 16.97 mm, *SE* = 2.45,P=1.38E-10). Finally, the FruitShapeIndexExternalI, i.e. the ratio of the maximum height to the maximum width (mA/mB), is increased by the minor allele (*β* = 0.91mA/mB, *SE* = 0.145, P=4.93E-09). Overall, the alleles of the SNP 3:183386147 specify whether a fruit is circular or elongated as measured by effect size on four phenotypes. The signal at the SNP 3:18338614 is the only one significant in region (±2 Mb) containing 26 genes (Figure 7a) and there is not high linkage disequilibrium with other genetic variants.

**Figure 7.**
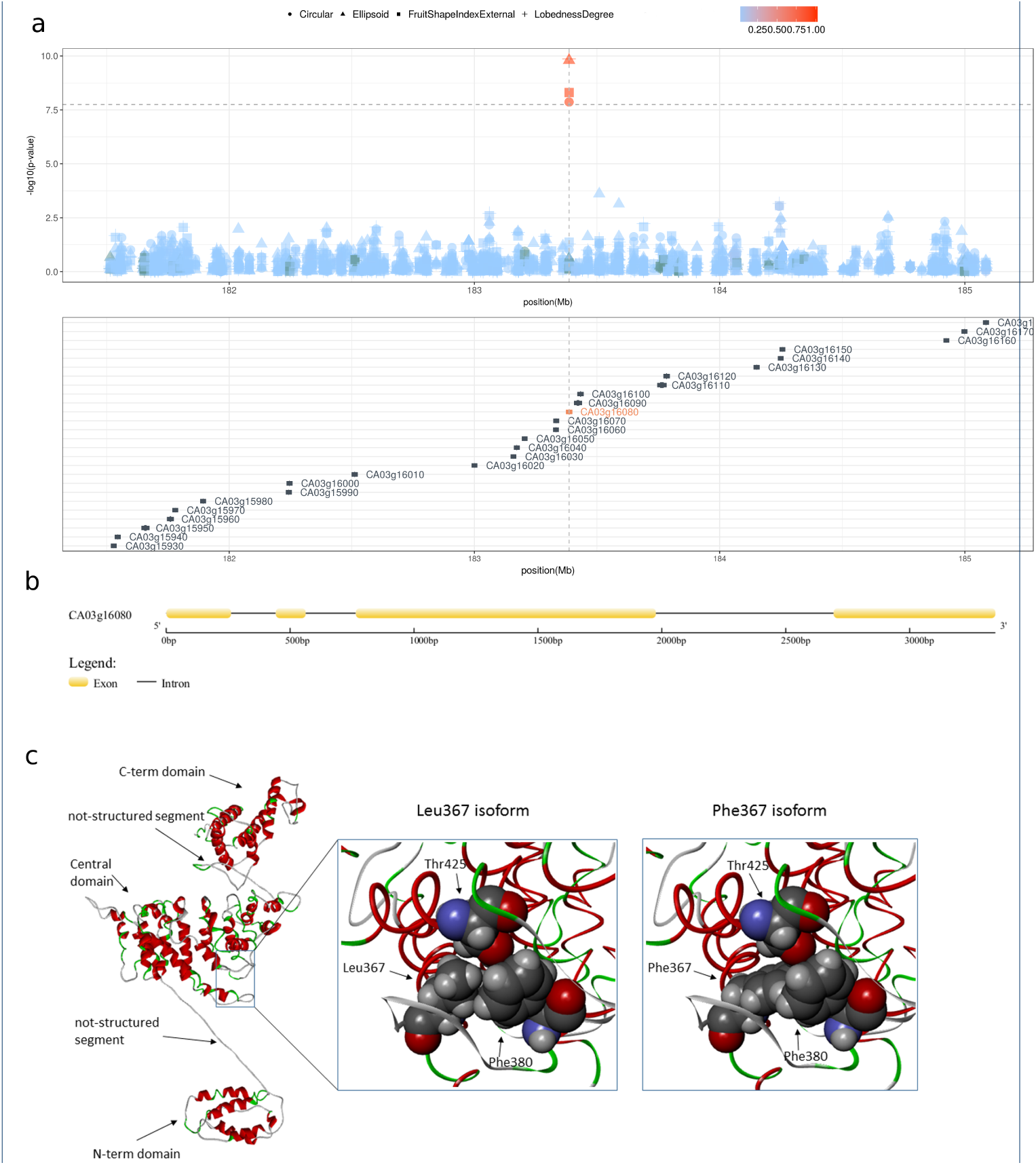
The Longifolia 1-like gene region. **(a)** Locus zoom plot in a region of ±2Mb surrounding the non-synonymous mutation (3:183386147) in the gene *Longifolia 1-like* (CA03g16080) showing that the 3:183386147 variant is the only one reaching a genome-wide significant threshold for genetic association in a region containing twenty-six genes. Color gradient indicate linkage disequilibrium measured ad r^2^. No other variants are in significant linkage with 3:183386147. **(b)** The predicted genic structure of *Longifolia 1-like* (CA03g16080). **(c)** Predicted protein structure for *Longifolia 1-like*. The whole model of the protein is represented by a backbone ribbon with helices in red and turns in green. Arrows indicate the structural regions, i.e. N-terminal domain, not structured connection segment, central domain, not structured connection segment, C-terminal domain. The region of the central domain including the residue number 367 (containing the 3:183386147 T→C variant) and the closer side chains (Phe380 and Thr425) is enlarged in spacefill representation in two versions, with the Leucine and the Phenylalanine residues. The representation with the Leucine highlights the compactness of the interactions between Leu367, Phe380 and Thr42)

We further investigated the population structure of *C. annuum* using PCA in order to perform PC-based selection scans[32] to find support for selection in the variants found in the GWAS. We used strict filtering on variants and samples to remove inter-species mixture and obtain a clean dataset of unadmixed *C. annuum* accessions to perform selections scans. The cleaned dataset consists of 212 *C. annuum* accessions and 100,773 variants. The principal components analysis is visualized in supplementary Supplementary Figure S7, where the first component mostly describes the genetic variance between European and Non-European samples and the second component captures the variance within the European samples. The first component is very likely to be associated with the ancestry component splitting the *C. annuum* accessions in the initial admixture plot from all species. The PC-based selection statistics of the top variants of the GWAS are reported in Supplementary Table S5 and Supplementary Figures S8 and S9 for most significant of the two first PCs. Only the 3:183386147 variant reaches nominal significance with a p-values of 0.015 and is thus not significant when taking multiple testing into account.

### A non-synonymous change in the *Longifolia 1-like* gene is associated with variance in *C. annuum* fruit elongation

The SNP 3:183386147 has two alleles, T and C. In our collection, the T allele is fixed in *C. pubescens, C. baccatum,* and *C. chacoense,* and nearly fixed (0.98) in *C. chinense* and *C. frutescens,* while its frequency in *C annuum* is 0.76, suggesting that T could be the ancestral state (Table 3). The SNP 3:183386147 is located in the third exon of the gene CA03g16080, which has no homologs in the pepper genome (Figure 7b). CA03g16080 is predicted to code for the protein Longifolia 1-like, a protein homologous to the Arabidopsis LONGIFOLIA1 and LONGIFOLIA2 and to *Oryza sativa* (rice) SLG7. Both LONGIFOLIA and SLG7 activate longitudinal organ expansion. [33, 34, 35] The mutation T→C causes a change in the aminoacidic composition from Phenylalanine to Leucine, with the removal of an aromatic group.

**Table 3.** Allele frequencies at the single nucleotide polymorphism 3:18338614 in *Capsicum* species. Numbers in parentheses indicate haploid sample size.

| Species (n) | CTC (Leu) | TTC (Phe) |
| --- | --- | --- |
| *C. annuum* (412) | 0.23 | 0.76 |
| *C. frutescens* (28) | 0.02 | 0.98 |
| *C. chinense* (90) | 0.03 | 0.97 |
| *C. baccatum* (72) | 0 | 1 |
| *C. chacoense* (26) | 0 | 1 |
| *C. pubescens* (26) | 0 | 1 |

We translated the DNA sequence of CA10g04730 into the corresponding protein sequence and predicted its secondary and tertiary structures to understand their structural properties and functional aspects. Functional terms associated to the protein indicate that it is a nuclear protein, as shown in Arabidopsis[33], with a possible involvement in protein binding and export from the nucleus, and RNA transport. The LONGIFOLIA 1-like protein contains 749 amino acids and is organized in a partial globular architecture, with three well-organized domains linked by segments without defined secondary structure (Figure 7c). The amino acid coded by the codon including the 3:183386147 T→C variant is the residue number 367 and is located at the edge of an *α*-helix of the central domain in a buried portion of the structure. The molecular modelling predictions indicate that the transition from phenylalanine to leucine in position 367 may cause a change in protein stability. The protein conformation with leucine seems more stable, probably due to the flexibility of its side chain, while the voluminous and rigid side chain of phenylalanine might induce a clash of side chains that might trigger a fold change, at least locally (Figure 7c).

## Discussion

*Capsicum* is one of the most extensively domesticated plants and its fruits are among the most-widely consumed. Although fruit morphology is a main target in breeding programs, the genetic basis of fruit shape has been studied so far by linkage analysis, using coarse sets of markers and low resolution phenotypic data. Similarly, the population structure of the *Capsicum* genus has never been investigated with a fine set of genetic markers. The recent availability of the reference sequence of a few species of *Capsicum* allowed the exploration of the genome properties, including the identification of genomic rearrangements among species. Nevertheless, little is known about genetic diversity within species due to the lack of studies with samples of adequate size. Our study is the first population study that combine sample size and depth of markers to highlight genomic features at species level and provide first insight into the genetics underlying morphological traits of pepper fruits.

The germplasm collection presented here covers all the economically important species of *Capsicum* widely used in breeding programs, representing the largest study so far in terms of number of species and number of genetic variants analyzed. Although limited to the 746k high quality variable sites accessible in the 1.8% of the genome, this study is an order of magnitude larger than previous studies in terms of the number of analyzed variants [23, 25], and also considers many more samples. The list of segregating sites positions and allele frequencies that we identified is publicly available and constitute a valuable resource for future studies.

We estimate that approximately 1.5% of *Capsicum* genome might be variable, therefore we expect that a number of variable sites in the order of 10^7^ might be discovered from population sequencing of the whole 3.6 Gbp of the genome. We discovered that genetic variants tend to form clusters since their average consecutive distance is lower than their average density. As expected genic regions are less variable than intergenic regions, suggesting greater selective constrains in genic regions.

Our work deciphers genetic variability at intra- and inter-species levels and associate it to morphological traits. We confirm and validate previous results showing that domesticated species tend to be less variable compared to wild ones[36] and that there is little admixture among species. We correlate to fruit size the major subdivision within *C. annuum* observed also in previous studies and never fully explained.[26, 2, 3] By determining that small and big fruits have different genetic backgrounds, we provide a rationale for genetic association studies for traits related to fruit morphology. In fact, we discover two significant associations. One links a clusters of genetic variants on chromosome 10 to traits determining whether a fruit is shaped or pointed. The other, a non-synonymous change on chromosome 3 in the gene *Longifolia 1-like,* is related fruit elongation and roundness. Both genomic regions overlap with QTLs linked to fruit shape and elongation previously identified in pepper[13, 12, 9, 18]. In particular major QTL detected on chromosome 3 (fs3.1) and confirmed in several studies (Supplentary Table 1) is located 10 Mbp upstream from the non-synonymous change in *Longifolia 1-like,* while a QTL on chromosome 10 (fs10.1) is located 100 Mb downstream the cluster of variants on chromosome 10. Those are certainly wide distances, probably related to the properties and limitations of QTL mapping, but it is encouraging to find overlap.

Among Solanaceae, the genetic basis of fruit shape have been extensively studied in tomato, leading to the identification of major QTLs and genes[7]. One of these, the Ovate-like gene CaOvate, has been reported in pepper to be involved in fruit elongation [37]. Beyond this, no other single genes information is reported, therefore, the genetic loci underlying fruit shape are not yet comprehensively explored. Despite the appropriateness of the promising association found in *Longifolia 1-Like,* further work will be necessary to fine map the signals that we found and fully understand the genetic asset behind pepper fruit shape and size.

## Conclusions

We present results form analyzing a small fraction of the *Capsicum* genome that are informative on overall genomic variability, population structure and genetic association that demonstrate the potential of making discoveries from population studies in this species. We can therefore predict that future studies based on more extensive sequencing and that can exploit haplotype-based information will allow both a better understanding of the evolutionary history of pepper (including reconstructing the domestication process and identifying footprints of natural selection), and discoveries that can guide the breeding process and inform genomic selection. As an example, a more in depth study of the properties of the non-synonymous change in *Longifolia 1-like* can guide applications in breeding that aim to modify pepper fruit appearance.

## Methods

### Genotype by sequencing, variant calling, and phasing

Genotype by sequencing was carried out as described in Taranto *et al.*[26]. Reads were mapped to the reference genome[22] using BWA [38]. Variant calling was done using Freebayes v1.2.0 [39] with standard parameters. Missing genotypes were imputed and the imputed genotypes were phased using Beagle v4.1 [40, 41].

### Population structure, phylogenetics, PCA

Model-based ancestry estimation was obtained using the ADMIXTURE software[42] with K ranging from 1 to 10. One thousand bootstrap replicates were run to estimate parameter standard errors. Ten-fold cross-validation (CV) procedure was performed and CV scores were used to determine the best K value.

### Plant material, growth conditions, phenotyping

Our world-wide collection of seeds from 373 accession belonging to the genus *Capsicum* includes both domesticated (landraces) and wild varieties (Table 1). Information about the accessions, including variety name, country of origin is in Supplementary Table S2. Three replicates for each accession were grown in a randomized complete-block design in greenhouse under controlled conditions at day-night temperature set points of 25/18 C. At maturity, eight fruits from each replicate were harvested, cleaned and cut longitudinally in two sections. Each section was scanned with a CanoScan LiDE 210 photo scanner (Canon, Tokyo, Japan) at a resolution of 300 dpi. Thirty-eight morphometric quantitative traits were recorded and analyzed using the Tomato Analyzer v 3.0 software[43]. A brief description of each trait, its acronym, and evaluation methodology are summarized in Supplementary Table S3 and visualized in Figure 5. Correlation between traits was calculated using the Spearman’s rank correlation test.

### Genome-wide association and PC-based selection scan

Genome-wide association analysis was performed using a linear mixed model implemented in GEMMA[44], controlling for admixture and relatedness between individuals as random effects using a genetic similarity matrix estimated from the data with the same software. Bonferroni-corrected genome-wide threshold for p-value is 1.786723e-08 and was calculated taking as reference a p-value of 0.01 and considering 559,684 markers. default filters in GEMMA.

We used PhonoGram[45] to display the chromosomal location along the pepper genome of the genome-wide signicantly associated SNPs.

PC-based selection scan was based on PC estimated using PCAngsd [46] as it is able to model the statistical uncertainty of genotypes and missing data and it has an implementation of the PC-based selection statistic of Galinsky et al. (2016)[32] working on genotype likelihoods. The filtering of variants was performed in ANGSD based on genotype likelihoods, and admixture analyses were performed in PCAngsd to filter out *C. annuum* samples which had an admixture proportion < 90% for the *C. annuum* cluster.

### Protein modelling and analysis

The exon-intron structure of the CA03g16080 gene was drawn using the Gene Structure Display Server 2.0. [47]

The amino acidic sequence of the gene CA03g16080 was used to generate a 3D model of the protein and to evaluate structural properties. In the absence of templates suitable for applying the homology modelling approach, we used a strategy based on the integration of predictions by means of different tools. We used PredictProtein[48] I-Tasser (Iterative Threading ASSEmbly Refinement) [49] for structure predictions and protein modelling. I-Tasser returned five best models and we selected three out of them, having the higher C-scores (from −1.55 to −1.85) and excluded the remaining two models, due to their lower C-scores (i.e., −4.03 and −4.62, respectively). The best models obtained have been used for evaluating structural properties by visual inspection with molecular viewers, and we integrated a complete evaluation with the results of other prediction tools, as and MAESTRO (Multi AgEnt STability pRedictiOn tool)[50], a tool for evaluating the possible effect of amino acid substitution on protein stability.

## Supporting information

## Ethics approval and consent to participate

Not applicable

## Consent for publication

Not applicable

## Availability of data and materials

Raw genetic data is available for collaboration upon request. Aggregate data (e.g. list of variable sites, allele frequencies, average phenotypic measures) are available on the GitHub repository https://github.com/ezcn/Capsicum-genomics

## Author's contributions

P.T. conceived and conducted the experiments. V.C., N.D.A., E.G., A.A. J.M., A.F., P.T. performed the analyses and worked at the interpretation of the results. V.C., N.D.A., E.G., P.T. wrote the manuscript. T.C. contributed to critically discuss the results. All authors reviewed and approved the manuscript.

## Acknowledgements

We are thankful to Roberto Sirica, Barbara Greco, and Francesca Taranto for valuable help in the first stage of the project. We also thanks Carmine Cirillo for contributing the git repository.

## Supplementary Figures

**Figure S1.**
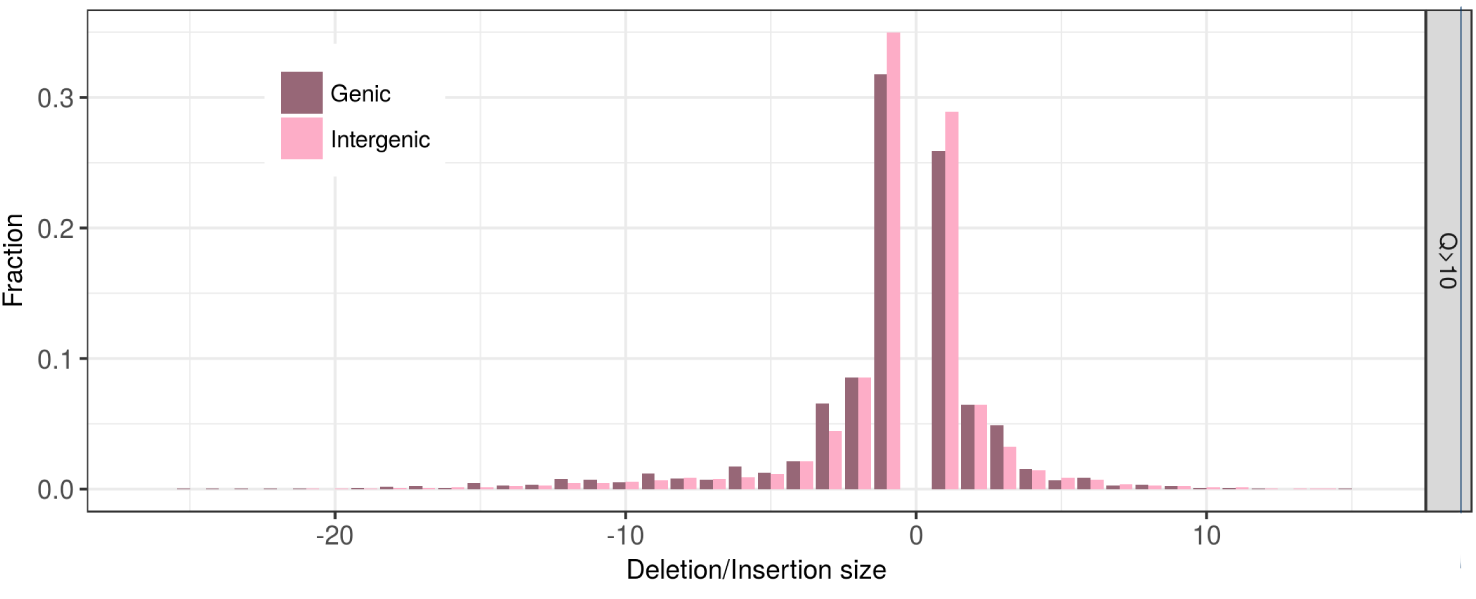
Size in nucleotides of deletions and insertion (InDel). Due to our use of GBS and reference-guided analysis, we were only able to discover InDels up to a few tens of bases. InDels of three or multiples of three nucleotides seems more frequent in genic region compared to intergenic ones, suggesting a preference for InDels that add or remove triplets over those causing frame-shifts mutations.

**Figure S2.**
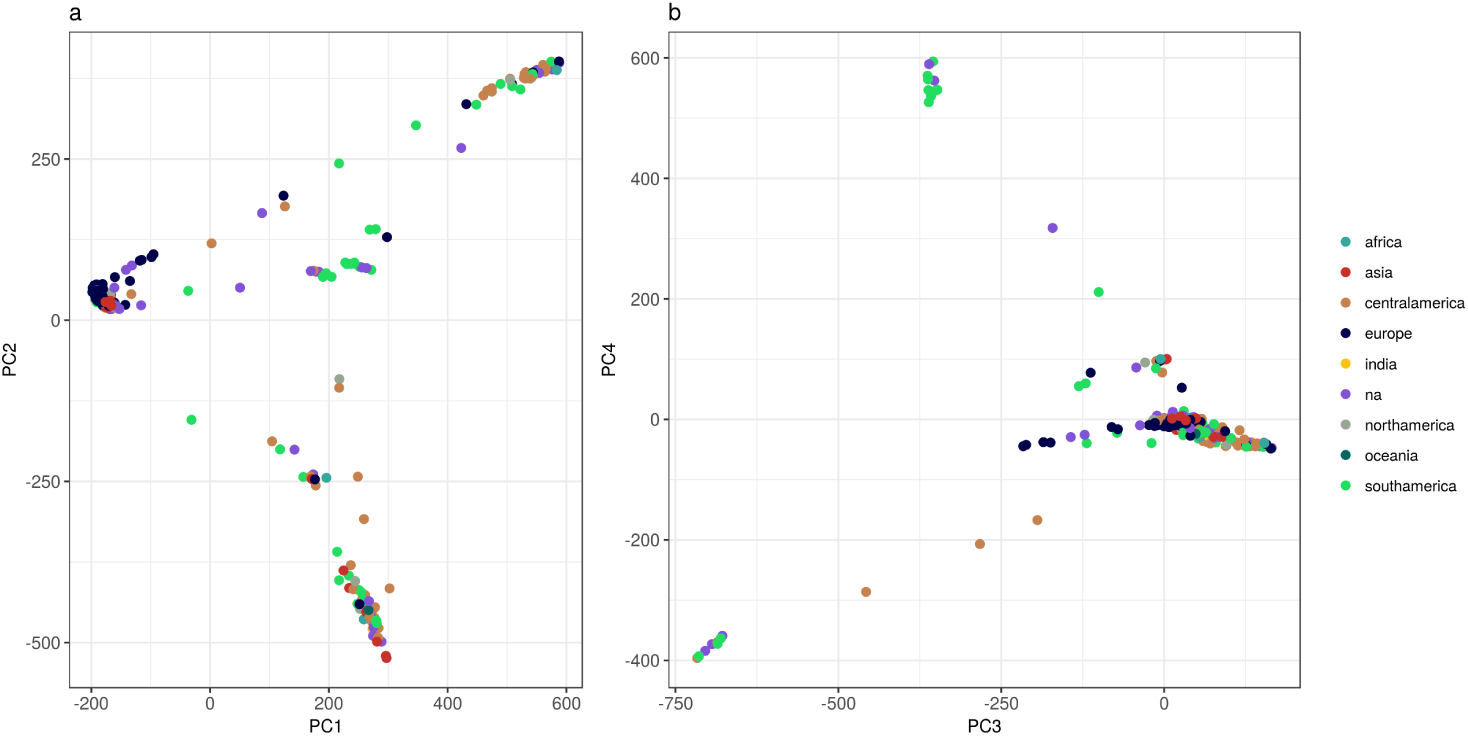
Principal component analysis (PCA) based on 746k genetic variants. Colors reflect the geographical origin of the accessions

**Figure S3.**
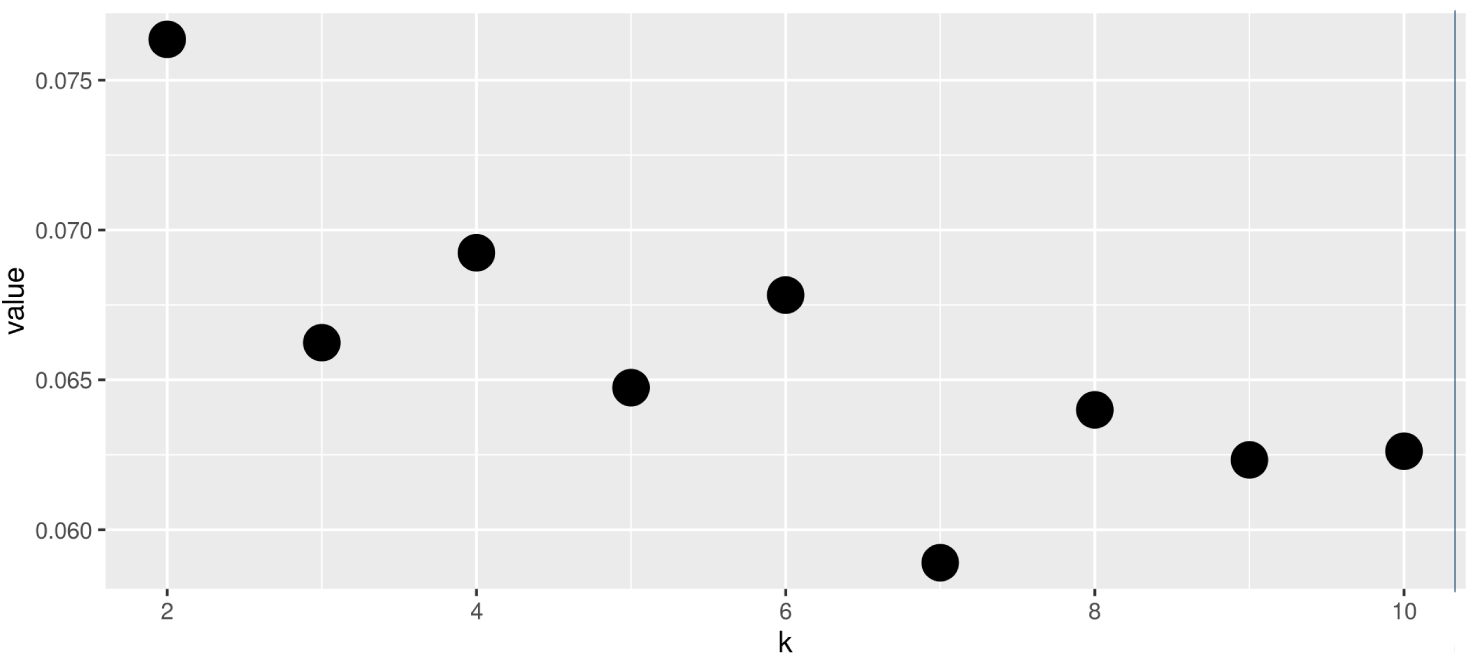
Cross-validation error of the admixture estimate for the hypotheses of 2 to 10 clusters.

**Figure S4.**
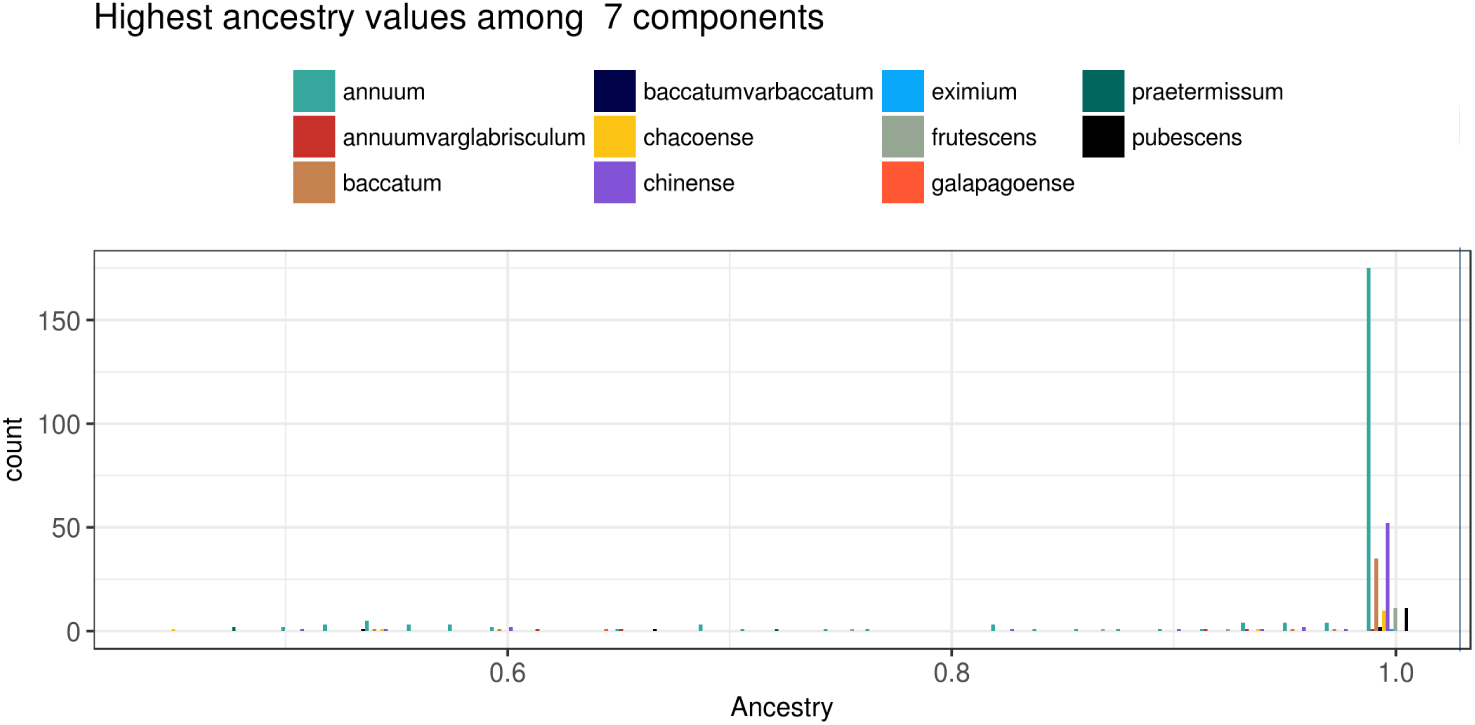
Admixture analysis. In the ADMIXTURE analysis most accessions belong to only one cluster, with the median coefficient of membership to the best-matching cluster being 0.99

**Figure S5.**
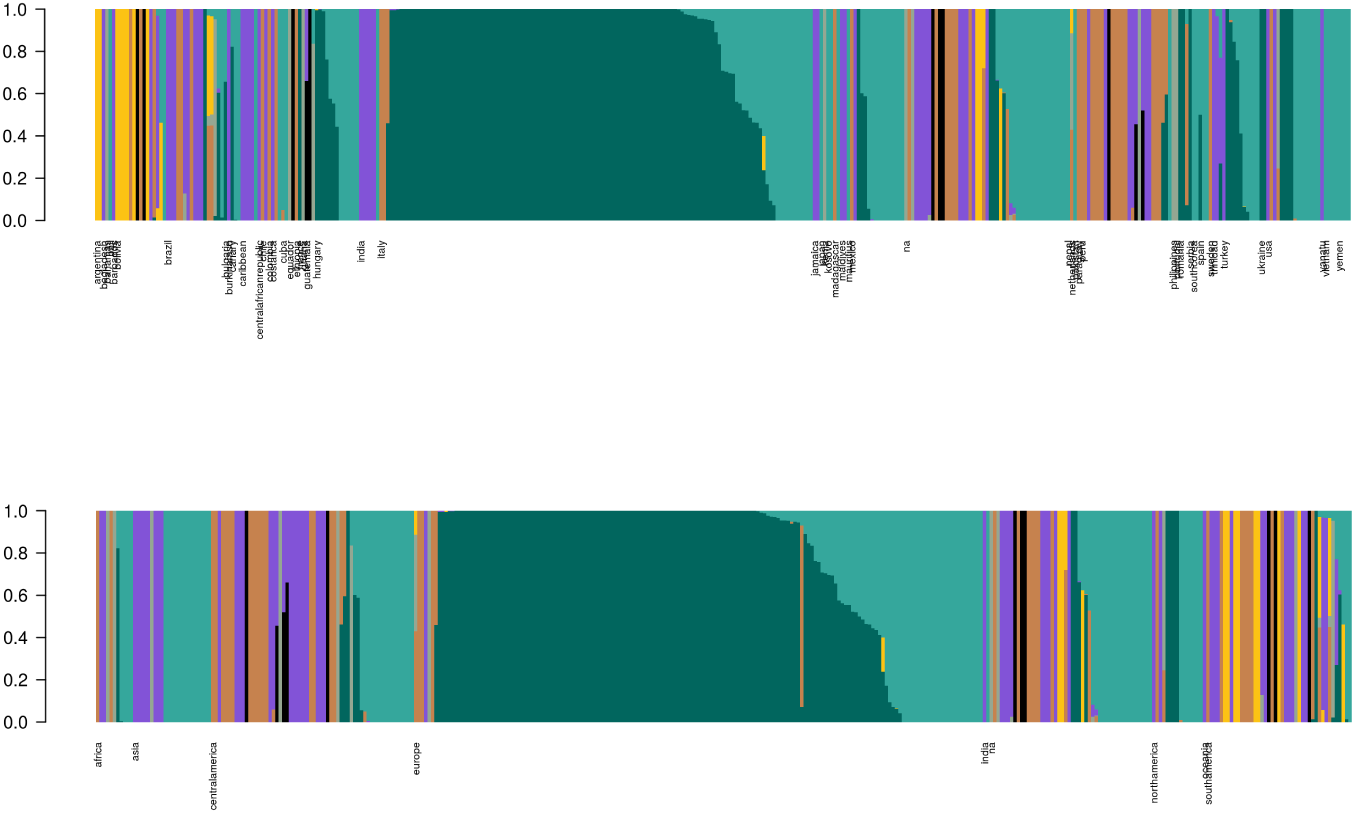
Admixture results in the hypothesis of 7 clusters. Accession have been grouped according to their geographical origin.

**Figure S6.**
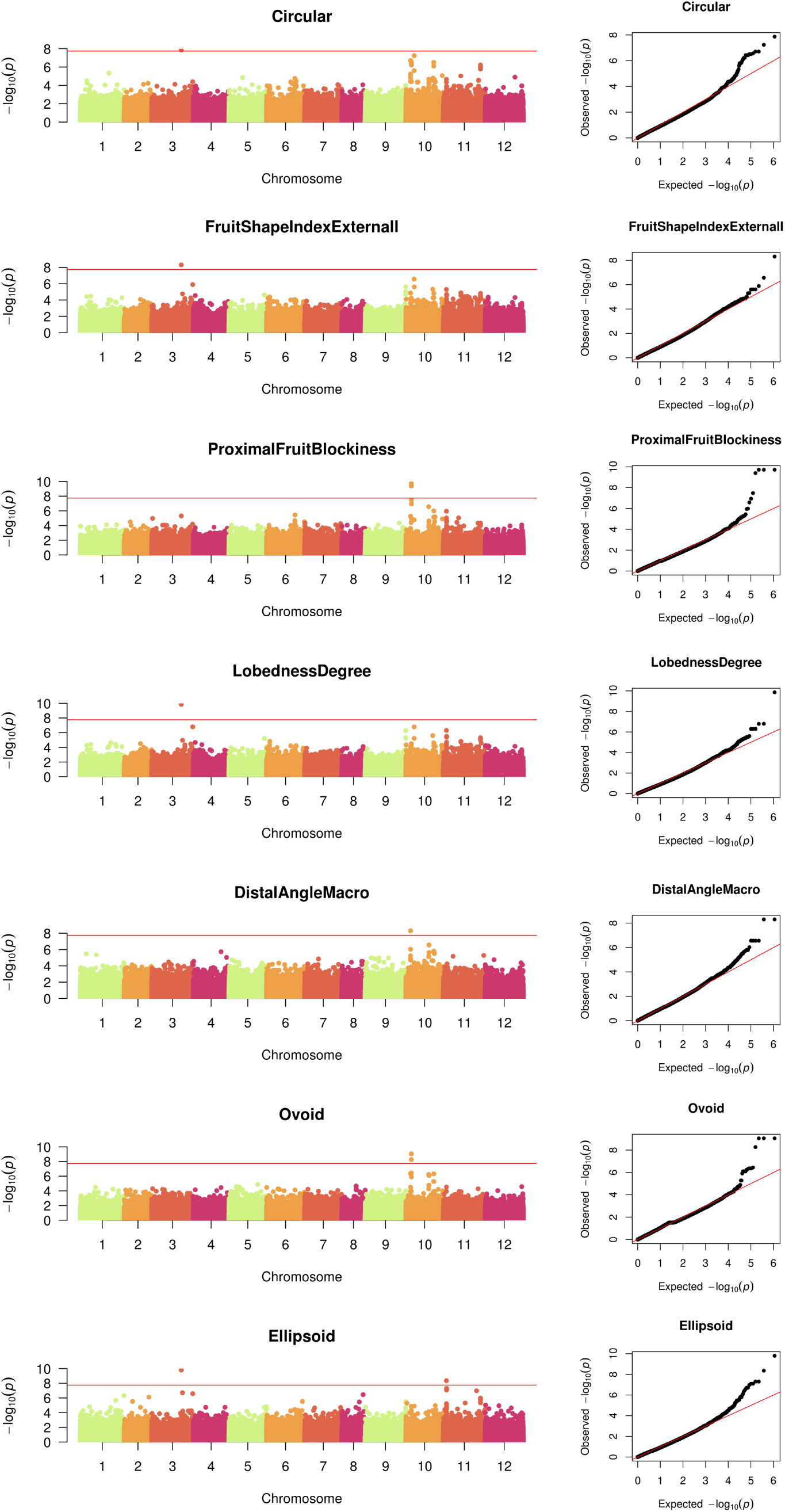
Manhattan plots and QQplots for the phenotypes presenting significant associations.

**Figure S7.**
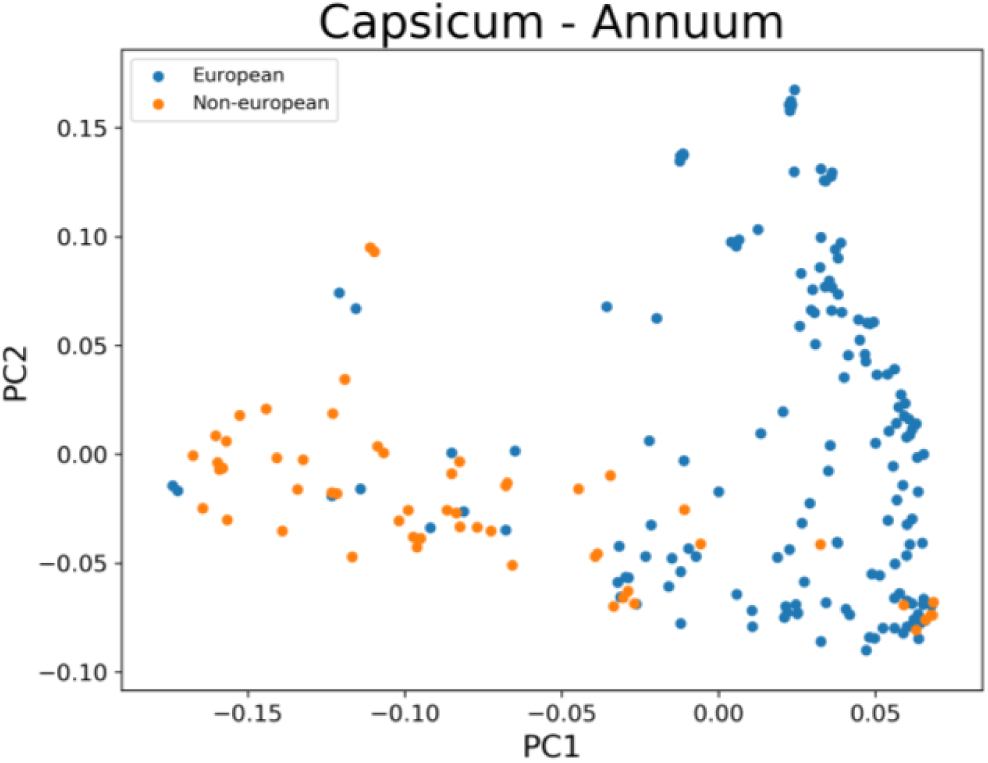
PCA plots of 212 unadmixed Annuum samples. The upper plot consists of the first and second PCs, and the bottom plot displays the first PC plotted against the third. PC1 visualizes the genetic variation between European and Non-european samples, PC2 visualizes the genetic variation in the European samples

**Figure S8.**
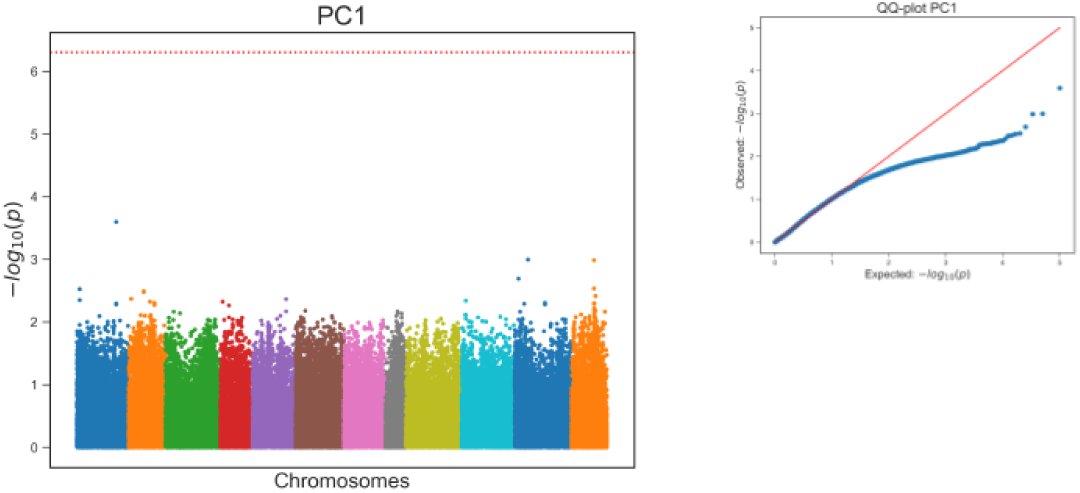
PC2 selection scan.

Additional file 2 — Supplementary Tables

